# Delineating the Effective Use of Self-Supervised Learning in Single-Cell Genomics

**DOI:** 10.1101/2024.02.16.580624

**Authors:** Till Richter, Mojtaba Bahrami, Yufan Xia, David S. Fischer, Fabian J. Theis

## Abstract

Self-supervised learning (SSL) has emerged as a powerful method for extracting meaningful representations from vast, unlabeled datasets, already transforming computer vision and natural language processing. Similarly, in single-cell genomics (SCG), representation learning is well-recognized for offering insights into complex biological data, even more so by the advent of early foundation model approaches. However, despite these advancements, identifying scenarios in SCG where SSL outperforms traditional supervised or unsupervised learning methods remains a nuanced challenge. Furthermore, selecting the most effective pretext tasks within the SSL framework for SCG is a critical yet unresolved question. Here, we address this gap by adapting and benchmarking SSL techniques in SCG, including masked autoencoders with multiple masking strategies and contrastive learning approaches. Trained on over 20 million cells, this study rigorously examines multiple downstream tasks, including cell type prediction, gene expression reconstruction, cross-modality prediction, and data integration. Our empirical analyses underscore the nuanced role of SSL, namely in transfer learning scenarios leveraging auxiliary data or analyzing novel datasets. Masked autoencoders excel over contrastive methods in SCG, diverging from computer vision trends. Moreover, our findings reveal notable capabilities of SSL in zero-shot cell type prediction and offer insights into its potential benefits in cross-modality prediction and data integration. In summary, we study the application of SSL in SCG, minimizing model bias through simple, fully connected networks, and benchmark SSL’s utility across key representation learning scenarios.

## Introduction

Single-cell genomics (SCG) has rapidly expanded into a big data domain, primarily driven by advancements in single-cell RNA-sequencing technologies^1^. This expansion has shifted the focus from analyzing data in isolated studies to using machine learning models for interpreting new data within the context of existing datasets^2^. Recent efforts towards comprehensive atlases, such as the Human Cell Atlas (HCA)^3^, underscore this development. However, larger datasets introduce new methodological challenges, such as technical batch effects across studies and the variability in labeling quality^4,5^. Large-scale models have gained interest and emerged for their potential to address these issues^6^. Yet a gap remains in understanding their use cases and how to effectively leverage the novel datasets comprising millions of cells^7^. The SCG field now not only requires computational power but also strategic use of methods that handle the complexities of big data. In this context, self-supervised learning (SSL) is a promising approach. SSL learns representations from data and pairwise relationships alone^8,9^, proven powerful in other data-intensive domains, such as computer vision^10,11^ and natural language processing^12,13^, leveraging large unlabeled datasets. It is thus often the basis for foundation models^14^.

SSL has already begun to impact SCG on small and large scales. On small scales, specialized SSL methods have deployed contrastive losses, tailored with techniques like multimodal learning^15^, graph-based strategies^16^, and clustering-based approaches^17–19^ to embed cells. The contrastive methods address unique data challenges in SCG, including batch effects and data sparsity^17,18,20–26^. Other specialized SSL methods predict blood cell traits^27^, identify sub-populations of T-cells^28^, boost active learning^29^, and classify cell types on the whole mouse brain^30^, indicating the method’s versatility. However, a common limitation among these approaches is their application to relatively small datasets or specific problems, resulting in limited generalizability across downstream tasks. On large scales, foundation models are trained on large datasets and applied to a broad range of tasks. In SCG, they often deploy transformers that are trained in a supervised^31,32^ and self-supervised^33–35^ fashion. While foundation models have demonstrated improvements through self-supervised pre-training^33,34^, disentangling the contributions of SSL, scaling laws, or the transformer architecture remains difficult. This ongoing debate underscores the relevance of investigating SSL in non-transformer contexts, which are prevalent in single-cell genomics^36,37^. Recent studies in computer vision^38,39^ also suggest a nuanced perspective on the dominance of transformer architectures, indicating the value of exploring diverse architectural approaches for model development. The ambiguity particularly arises when comparing the performance of models with and without self-supervised pre-training^13,40^, suggesting a need for a more in-depth exploration of SSL’s role in the evolving landscape of SCG.

To guide the effective usage of SSL in SCG, we need to address these ambiguities through systematic empirical validation. Such a study helps to determine the scenarios in which SSL can effectively contribute to SCG. First, this requires developing SSL methods based on first principles and tailoring them for single-cell applications. These SSL methods leverage pairwise relationships within data *X* for training, setting them apart from supervised learning, which relies on data *X* with labels *Y* to guide the loss, and unsupervised learning, which depends solely on data *X*^8,9,41^. Subsequently, we benchmark our SSL methods and compare their performance to their supervised and unsupervised counterparts. Second, this study requires validation across downstream applications, addressing the method’s objective to learn data representations that are helpful across multiple tasks.

Our study aims to identify specific scenarios in SCG where SSL is helpful and to thoroughly analyze and evaluate SSL approaches in SCG. Utilizing the scTab dataset^4^, which comprises over 20 million cells, our study assesses the effectiveness of SSL across multiple downstream tasks. Based on well-defined benchmark metrics for SSL in SCG, our empirical analysis primarily focuses on the cell-type prediction application, with validation in gene expression reconstruction, cross-modality prediction, and data integration. We find that SSL improves downstream performance in transfer learning settings, that is, when analyzing smaller datasets informed by insights from a larger auxiliary dataset and in scenarios involving novel, unseen datasets. This improvement is especially notable in class-imbalance-sensitive metrics, indicating robustness improvements. However, our findings also reveal that self-supervised pre-training on the same dataset as the fine-tuning does not yield improvement compared to only supervised or unsupervised training. In summary, our study clarifies the roles and benefits of SSL in SCG, demonstrating its strengths in specific contexts while identifying its applicability limits. This research contributes to a more informed and strategic use of SSL in SCG, particularly in advancing our understanding of complex biological datasets.

## Results

### A Self-Supervised Learning Framework for Single-Cell Genomics

We present an SSL framework to develop self-supervision methods and study different use cases in SCG. Central to our framework is the use of fully connected autoencoder architectures, selected for their ubiquitous application in SCG tasks^36,37^ and for minimizing architectural influences on our study, yet still large enough to capture underlying biological variations. In this framework, we integrate key SSL pretext tasks based on masked autoencoders^42^ and contrastive learning^43,44^ to benchmark their performance. The framework operates in two stages: The first stage is pre-training, also called pretext task, where the model learns from unlabeled data. We call the resulting model ‘SSL-zero-shot’ for its zero-shot evaluation. The second stage is the optional fine-tuning. We call the resulting model the ‘SSL’ model, which is further trained to specific downstream tasks such as cell type annotation (see Figure 1a). The pretext task builds a rich data representation based on a comprehensive dataset. We chose the scTab dataset^4^ because of its extent and diversity. We used all 19,331 human protein-encoding genes from scTab to maximize generalizability, ensuring gene coverage for analyses of novel datasets, regardless of their feature selections. Our SSL framework leverages Masked Autoencoder with Random Masking and Gene Program Masking (GP) strategies, along with our Isolated Masked Autoencoder (iMAE) approaches Gene Program to Gene Program (GP to GP) and Gene Program to Transcription Factor (GP to TF) masking, considering isolated sets of genes (see Figure 1b). The strategies entail leveraging different degrees of biological insight, from random masking with a minimal inductive bias to isolated masking that intensively utilizes known gene functions, emphasizing targeted biological relationships. For contrastive learning, we incorporate Bootstrap Your Own Latent (BYOL)^43^ and Barlow Twins^44^, known for their effectiveness in computer vision (see Figure 1c). We benchmarked these strategies for their efficacy in improving downstream performance. Our SSL framework, including these strategies, is depicted in Figure 1a, outlining its architecture and pivotal components. Detailed descriptions of the specific implementations and adaptations of these SSL methods for SCG are further elaborated in the Methods section.

**Figure 1:**
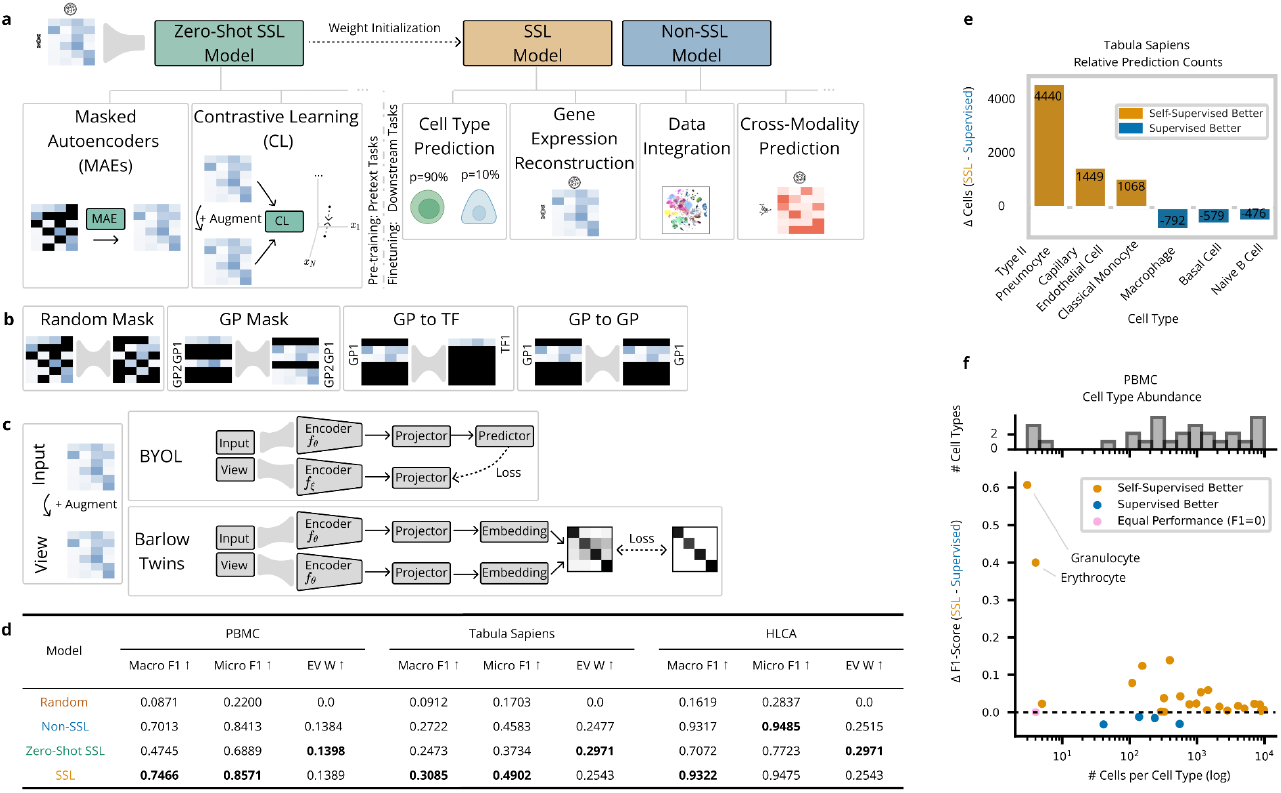
Self-supervised learning (SSL) on auxiliary data in Single-Cell Genomics (SCG) improves downstream performance. **(a)** Overview of the SSL framework used in this paper (Methods). The Zero-Shot SSL model is trained with self-supervision algorithms, Masked Autoencoders, and Contrastive Learning on RNA-sequencing data from scTab. Its weights are used as initialization for the SSL model, which is then fine-tuned for the downstream task, such as cell type prediction or gene expression reconstruction. The Non-SSL model is initialized randomly and only fine-tuned for the downstream task. **(b)** Masking strategies: Input features are zeroed out (black), and non-black features are unchanged. The autoencoder (grey) predicts the features are on the right side. Only the non-black predicted features are used for the loss. Further depicted are gene programs (GPs) and Transcription Factors (TFs) (Methods). **(c)** Contrastive learning methods. The Input is augmented with a data augmentation to the View. Bootstrap Your Own Latent (BYOL) and Barlow Twins are contrastive models to obtain the data representation (Methods). **(d)** Result of the smaller datasets experiments. Evaluated are (i) a random model or random performance, (ii) a Zero-Shot SSL model, (iii) a Non-SSL model (that is, *e*.*g*., supervised for cell type prediction and unsupervised for gene expression reconstruction), and (iv) an SSL model. The models are deployed on three datasets: PBMC, Tabula Sapiens, and HLCA (Methods). Cell type prediction is evaluated with the macro and micro F1 score (higher=better), and the gene expression reconstruction is evaluated with the weighted explained variance (EV W, higher=better). **(e)** Relative number of cells correctly predicted by the SSL model and the supervised model plotted for the cell types with the largest absolute performance difference. **(f)** macro F1-score differences for the SSL model and the supervised model plotted against the number of cells per cell type, *i*.*e*., the abundance of cell types. Additionally, the number of cell types per abundance is plotted above.

### Self-supervised pre-training on auxiliary data improves cell type prediction

As a first use case for self-supervision in SCG, we asked if analyses on cell atlases or smaller datasets can benefit from self-supervised pre-training on auxiliary data. We answered this using three datasets: the Human Lung Cell Atlas (HLCA)^5^ (2,282,447 cells, 51 cell types), Peripheral Blood Mononuclear Cells (PBMCs) after SARS-CoV-2 infection^45^ (422,220 cells, 30 cell types), and the Tabula Sapiens Atlas (483,152 cells, 161 cell types)^46^. These datasets vary in size, biological context, and complexity, providing a robust test bed for our models. We evaluated cell type prediction with the macro F1 score, supplemented by the micro F1 score, to compare robustness against class imbalances. We evaluated gene expression reconstruction with the Mean Squared Error (MSE). For the PBMC and Tabula Sapiens datasets, the self-supervised pre-training on additional scTab data significantly improved cell type prediction and gene expression reconstruction (see Figure 1b and Supp Figure 2): From [0.7013 ± 0.0077] to [0.7466 ± 0.0057] macro F1 in the PBMC dataset and from [0.8413 ± 0.0034] to [0.8571 ± 0.0017] macro F1 in the Tabula Sapiens dataset. In the Tabula Sapiens dataset, this improvement is driven by strongly enhancing the classification of specific cell types, correctly classifying 6881 of 7717 Type II Pneumocytes instead of 2441 (see Figure 1e). For the PBMC dataset, this improvement is pronounced for underrepresented cell types (see Figure 1f), also indicated by the stronger macro F1 improvement versus micro F1 improvement. In contrast, the HLCA dataset presented a marginal performance improvement through self-supervised pre-training.

### A tailored pre-training strategy leads to high zero-shot performance

The scenario in which SSL is typically evaluated in computer vision is the zero-shot setting, where the model’s ability to represent and distinguish unobserved classes is assessed using data representations obtained solely through self-supervised pre-training. The labels are predicted, *e*.*g*., with kNN classification or by training a prediction head while freezing the encoder weights. This perspective is noteworthy in SCG, where datasets’ increasing volume and complexity often come with challenges in obtaining accurate and comprehensive labels^5^. The ability of Zero-Shot SSL to achieve up to a 0.6725 macro F1 score on the scTab test set stands out as a strong performance (see Figure 2a). Likewise, in the test cases of HLCA, PBMC, and Tabula Sapiens, Zero-Shot SSL comes close to their fine-tuned counterparts (see Figure 1d, Supp Figure 1). The embedding from the zero-shot model illustrates this implicitly learned distinction of cell types (see Figure 2b). These findings highlight SSL’s potential in SCG to reduce the reliance on curated labels^47^ and propose adding self-supervised pre-trained model embeddings to biological analyses alongside PCA, a practice exemplified by platforms like CELLxGENE^48^. However, our benchmarking of SSL methods revealed the sensitivity to the choice of pre-training strategy. While contrastive methods have shown efficacy in other domains^41,43,44^ and specialized in smaller scales in SCG^17,18,20–26^, our study found that standard contrastive approaches did not yield as promising results for diverse, large-scale SCG tasks. This result highlights the challenges of applying these methods as generalizable pretext tasks for single-cell data. Conversely, Masked Autoencoders performed better: the random masking strategy consistently ranked among the top performers across different tasks (see Figure 2a). Notably, in the specific context of gene expression reconstruction, the Gene Program to Transcription Factor (GP to TF) isolated masking demonstrated superior performance compared to other methods (See Supp Figure 1 and 2). This finding highlights the potential of tailored masking strategies in capturing the nuanced biological variations inherent in SCG data.

**Figure 2:**
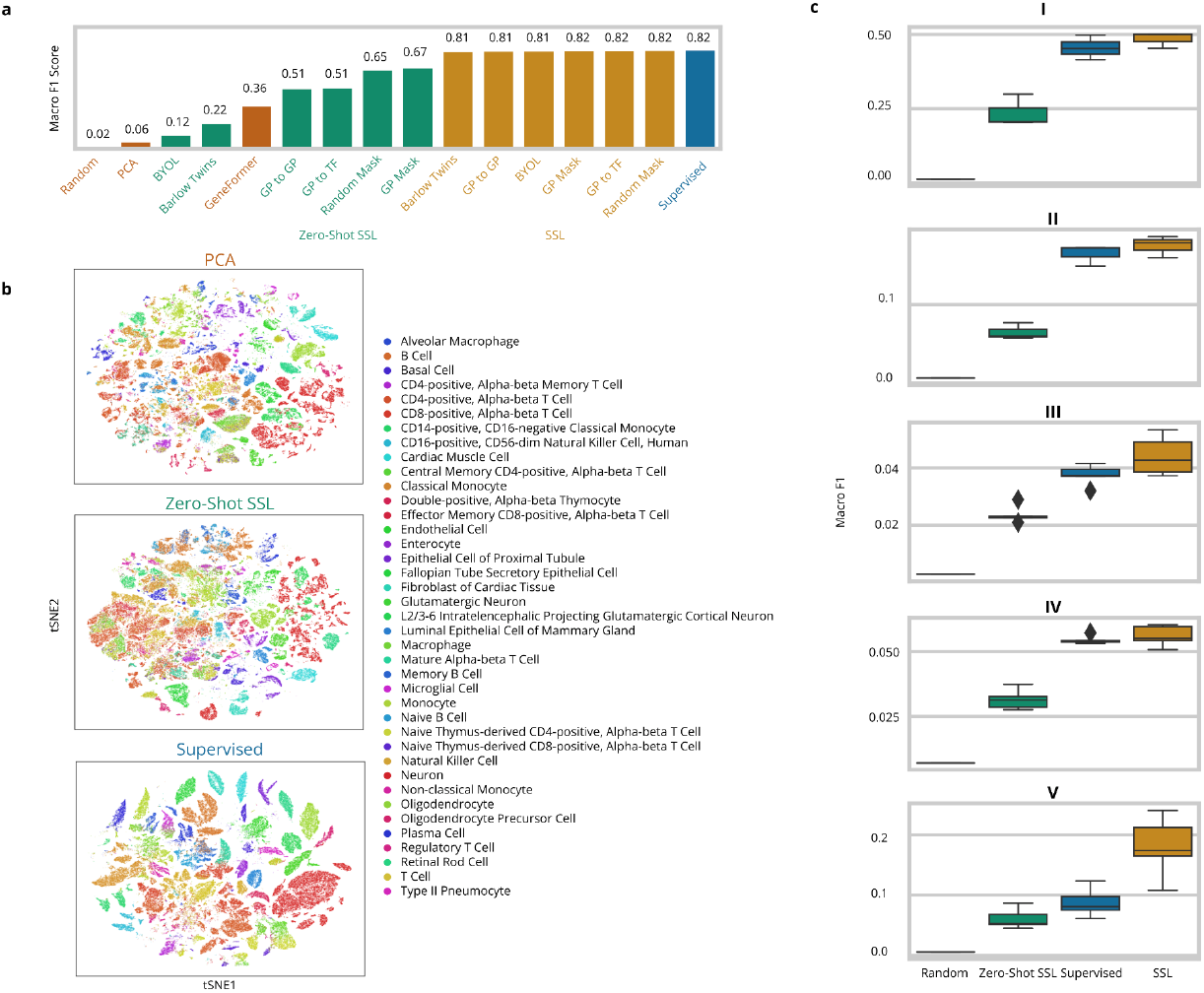
SSL enables high zero-shot performance and higher accuracy on unseen datasets. **(a)** Benchmark result of cell type prediction on the scTab holdout test set using kNN classification (Methods). We compare (i) baseline methods of kNN classification on a randomly initialized model, on the PCA embeddings, and deploying GeneFormer^33^ in a zero-shot setting; (ii) our Zero-Shot SSL methods, pre-trained on the scTab training data; (iii) our SSL methods, and (iv) the supervised model. **(b)** t-SNE visualization of the baseline PCA embedding, the embedding obtained from the Zero-Shot SSL model, and the supervised model. **(c)** Classification performance on unseen datasets (I: Human Brain Atlas - Tail of Hippocampus (HiT) Caudal Hippocampus -CA4-DGC.^49^ II: Human Brain Atlas All non-neuronal cells.^49^ III: Single-cell analysis of prenatal and postnatal human cortical development.^50^ IV: Circulating Immune cells after CV19 infection, vaccination, and HC.^51^ V: Human: Great apes study^52^) measuring the macro F1 Score of a random baseline, a Zero-Shot SSL model, the supervised model, and an SSL model.

### The efficacy of SSL depends on its context

While the prior evaluations focused on carefully curated and widely used benchmarks, we also set out to investigate SSL’s nuanced behavior in analyzing in-distribution versus novel data settings. If the supervised and SSL model are provided access to the same data, their performance is remarkably similar (see Figure 2a). This finding is notable across cell type annotation and gene expression reconstruction. Extending to novel datasets, we evaluated the supervised and SSL models on five datasets^49–52^ published after the CELLxGENE^48^ census of scTab (Methods). In this setting, self-supervised pretraining improves performance (see Figure 2c, Supp Figure 2), *e*.*g*., from [0.0877 ± 0.0215] to [0.1797 ± 0.0450] macro F1 for cell type prediction in the Great Apes study^52^. So, while inside the distribution (Figure 2a), supervised and self-supervised learning perform similarly, this finding offers another dimension of SSL’s utility: when analyzing novel, unseen datasets, where generalization is crucial, SSL shows its advantages.

### Improving Cross-Modality Prediction Performance through SSL Pre-training

Having benchmarked the utility of SSL on transcriptomics, we extended our study to multiomics^53^, posing the question: can SSL leverage auxiliary data from one modality to enhance multimodal downstream tasks, here focusing on cross-modality prediction (see Figure 3a). The NeurIPS multiome dataset^54^, a rich multi-donor, multi-site, and multi-modal bone marrow dataset containing coupled gene expression and proteomics counts from CITE-seq^55^ experiments, provided a suitable test bed. The models obtain RNA-sequencing counts as input and predict protein counts. The SSL models are additionally pre-trained on RNA-sequencing data from the auxiliary scTab and the NeurIPS multiome dataset. When pre-trained on scTab, SSL significantly outperforms its supervised counterpart and the baseline method totalVI^56^ (see Figure 3b,c). The Pearson correlation between predicted and true protein counts improved from [0.8627 ± 0.0026] for the supervised model to [0.8713 ± 0.0012] for the self-supervised model. Notably, the improvement is smaller if pre-trained on the same data, to a Pearson correlation of [0.8654 ± 0.0024]. This finding highlights the advantage of self-supervision in cases where one modality is more abundant.

**Figure 3:**
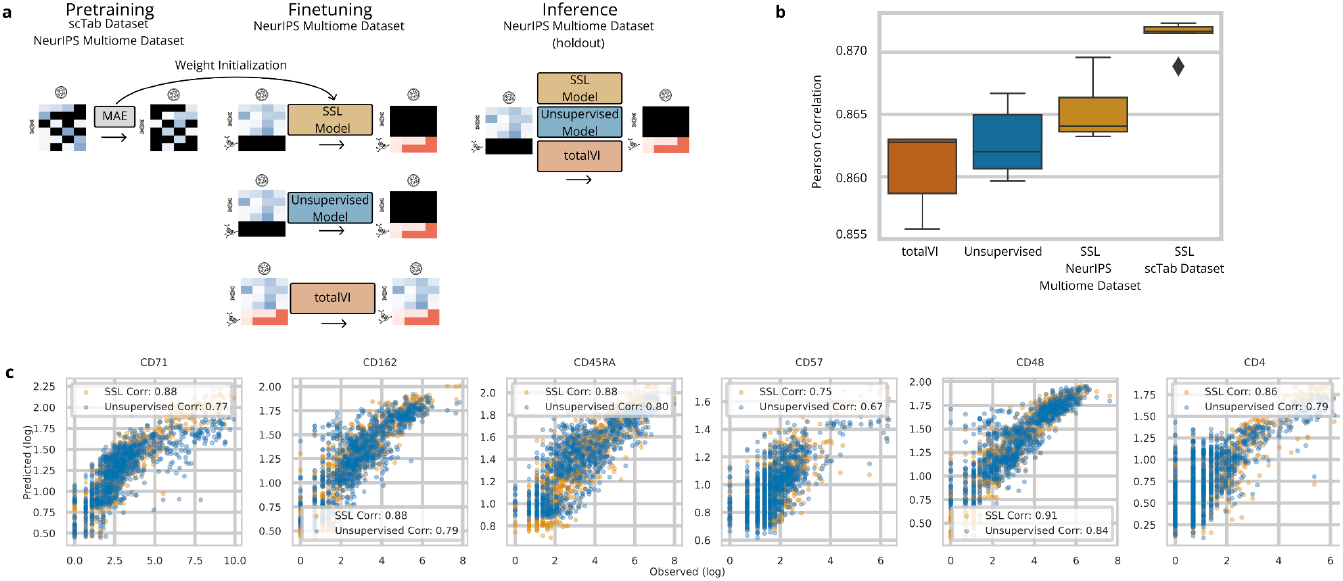
Self-supervised pre-training on auxiliary data improves cross-modality prediction. **(a)** Scheme of cross-modality prediction training. The SSL models are pre-trained with Masked Autoencoders (MAE) on RNA-sequencing data (i) of the downstream NeurIPS multiome dataset or (ii) of the auxiliary scTab dataset. The SSL model, initialized with the pre-trained weights, and the supervised model, randomly initialized, predict the protein counts from the RNA counts. The baseline totalVI model learns a joint distribution of RNA and protein counts. In inference, all models predict the protein counts from a holdout test set. **(b)** Cross-modality prediction performance, Pearson correlation between predicted and true protein counts. The linear regression baseline with a Pearson correlation of 0.8085 is not shown for visualization purposes. **(c)** Scatter-plot of predicted log counts against true log counts for exemplary proteins.

### Self-supervised pre-training enhances data integration

Integrating single-cell datasets for joint analysis is difficult due to batch effects, *e*.*g*., experimental conditions or confounding factors, posing unique challenges to atlasing efforts^5^. Large-scale models in SCG have already been deployed to address this challenge^31,33^. To clarify the role of SSL in these efforts, we set out to integrate three datasets included in scTab: the molecular cell atlas of the human lung (65,662 cells, 45 cell types)^57^, the molecular atlas of lung development of LungMap (46,500 cells, 28 cell types)^58^, and the molecular single-cell lung atlas of lethal COVID-19 (116,313 cells, 30 cell types)^59^. The datasets vary in cell type composition and donor health or disease states, providing a challenging environment for this task. The scIB metrics^60^ evaluate data integration performance, indicating how well batch effects are corrected while conserving the biological variability (see Figure 4a). We fine-tuned the supervised and SSL models using gene expression reconstruction on these three datasets. To improve data integration performance and model comparison, we added batch covariates to all models^53^. This led to the SSL shallow model, which fine-tunes the last encoder layer of the Zero-Shot SSL model with batch covariates. PCA and scVI^36^ embeddings serve as baselines for data integration. The scIB metrics indicate that self-supervised pre-training improves the data integration performance (see Figure 4b) with a total scIB score of 0.57 (SSL-shallow) and 0.55 (SSL) compared to 0.52 (supervised). The SSL-shallow model performed best, hinting at a meaningful data representation learned through the self-supervision algorithms, underscored by the comparable performance of the specialized data integration method scVI^36^. This finding supports the advantage of leveraging auxiliary data through self-supervised learning and showcases the effectiveness of minimal fine-tuning compared to supervised learning.

**Figure 4:**
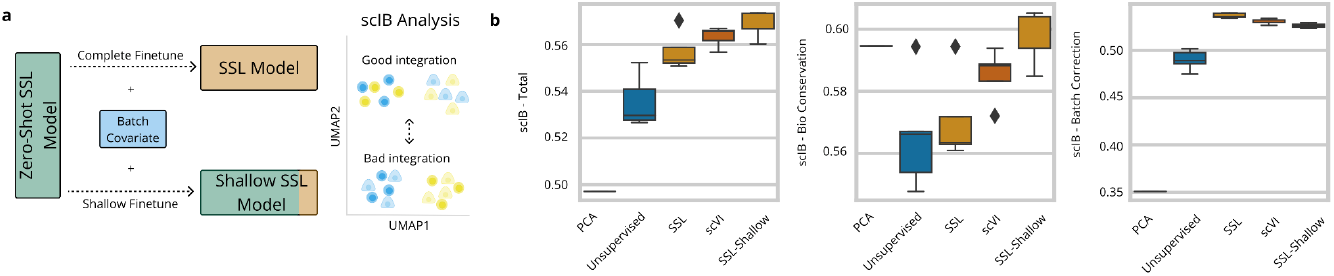
Self-supervised learning benefits data integration. **(a)** Scheme of data integration fine-tuning for SSL and SSL-Shallow models. The SSL-Zero-Shot model is fine-tuned, along with the batch covariates available for the data integration benchmarking. A complete fine-tuning results in an SSL model, and a shallow fine-tuning of the last encoder layer results in an SSL-Shallow model for the data integration task. **(b)** The scIB data integration benchmarking results^60^ of 5 runs. The benchmarking analysis comprises two sets of bio conservation and batch correction scores aggregated into a total score with a weighted mean.

## Discussion

We analyzed the application of self-supervised learning in single-cell genomics to guide its effective usage, leading us to adapt and benchmark several SSL techniques tailored for SCG. Our empirical study illuminates the context in which SSL can excel, especially when leveraging insights from vast, auxiliary datasets for smaller dataset tasks and in novel dataset scenarios. We also demonstrated that SSL shows parity with supervised methods where both access the same data, and that the Zero-Shot SSL model comes close to that performance. Our insights contribute to a more nuanced understanding of SSL’s applications in SCG. By rigorously testing these methods on an expansive dataset encompassing over 20 million cells, we offer a robust, empirically grounded perspective on the use of SSL in SCG, paving the way for more informed, data-driven approaches in studying complex biological systems. In the context of large-scale and foundation models^33–35^, this understanding could help design pre-training and select pretext tasks. For broad applicability within SCG we address diverse, meaningful tasks from cell type prediction, gene expression reconstruction, cross-modality prediction, and data integration. By demonstrating that SSL’s advantages emerge predominantly in scenarios involving transfer learning tasks through auxiliary data or distributional shifts, we offer a pragmatic lens through which the SCG community can view SSL - not as a universal solution but as a strategic tool tailored for specific challenges. This insight is particularly relevant as SCG moves towards larger data analyses, analyzing cell atlases and leveraging the consortia of millions of cells in foundation models. The adaptability and robustness of SSL, as evidenced in our empirical analysis, are crucial in this context to leverage large datasets. Our approach thus shows an example in SCG for the contextual application of SSL, guiding researchers to leverage this methodology where it most effectively addresses the field’s unique data challenges.

Future work on SSL in SCG may follow up on our findings. First, we identified several scenarios where SSL can improve performance across downstream tasks. These scenarios can serve as a baseline for future work, such as adding further downstream applications or developing novel SSL methods. Second, we benchmarked several SSL methods tailored for SCG. While Masked Autoencoders (MAEs) achieve high performances, it is notable how the performance differs by masking strategies and how contrastive learning has not yet achieved its full potential. We believe it is due to the default data augmentations. Finding a tailored yet generalizable data augmentation for contrastive learning in SCG remains an interesting challenge. Our framework for SSL evaluation across tasks and scenarios can serve as a baseline. Third, the remarkable performance improvement through SSL pre-training on auxiliary data promises further applications in which the data or data modality is scarce. In applications such as dynamics modeling, where datasets with temporal resolutions are limited in size and availability, or in smaller applications where the dataset size is very small, our solution can potentially improve analysis performance.

Finally, our study clarifies the scenarios in which SSL pre-training can improve performance in SCG. Namely, SSL excels in transfer learning tasks through leveraging auxiliary data and in distributional shift scenarios. In the context of foundation models, we illuminate methodological innovations stemming from the SSL pre-training. For the broader computational biology community, we have shown that self-supervised pretraining on atlas-level data can help to improve performance on smaller datasets of biological or medical relevance that are commonly more difficult to scale.

## Methods

### Data Curation

#### Preprocessing

All datasets employed in this study underwent a commonly used preprocessing pipeline in single-cell genomics. This involved normalization to 10,000 counts per cell and log1p transformation to mitigate technical variations and facilitate more meaningful biological comparisons. This uniform preprocessing approach ensured that our models were trained and evaluated on data closely reflecting the underlying biological realities while minimizing technical noise. *scTab Dataset*

The core dataset for our study stems from scTab^4^ and is derived from the CELLxGENE^48^ census version 2023-05-15, a long-term supported release hosted by CELLxGENE. This dataset represents a substantial collection of human single-cell RNA-sequencing data, encompassing 22.2 million cells spanning 164 unique cell types, 5,052 unique donors, and 56 different tissues. To ensure the reproducibility of dataset creation, scTab applied stringent criteria for inclusion, focusing on primary data from 10x-based sequencing protocols and ensuring a broad representation across cell types and donors. The scTab data is divided into training, validation, and test sets based on donors, avoiding label leakage and ensuring each set contains unique donors. This donor-based splitting approach allowed us to maintain a proportional representation of cells across the sets. It ensured that each cell type was represented in the training and testing phases. It further presented a challenging test split with unseen donors. The final split resulted in 15.2 million cells for training, 3.5 million for validation, and 3.4 million for testing.

#### Single-Cell Atlases

We further considered smaller, focused datasets to test if access to the auxiliary data gives an advantage. These datasets are subsets of the scTab data, tailored to specific applications and consist of the Human Lung Cell Atlas (HLCA)^5^ (2,282,447 cells, 51 cell types, 540,732 train, 117,541 validation, 117,517 test samples), Peripheral Blood Mononuclear Cells (PBMCs) after SARS-CoV-2 infection^45^ (422,220 cells, 30 cell types, 78,354 train, 33,761 validation, 189,756 test samples), and the Tabula Sapiens Atlas (483,152 cells, 161 cell types, 223,337 train, 54,908 validation, 57,616 test samples)^46^. The division into train, validation, and test sets is derived from their allocation within the scTab dataset to prevent data leakage.

#### Novel Datasets

To evaluate our models’ performance in novel data analysis scenarios, we incorporated five unseen datasets published after the CELLxGENE census version of scTab: (i) All non-neuronal cells from the Human Brain Atlas^49^ (ii) Dissection: Tail of Hippocampus (HiT) - Caudal Hippocampus - CA4-DGC from the Human Brain Atlas^49^, (iii) the Single-cell analysis of prenatal and postnatal human cortical development^50^, (iv) Circulating Immune cells -- CV19 infection, vaccination and HC^51^, and (v) Human: Great apes study^52^. The novel datasets were filtered for the genes used in scTab; missing genes were zero-padded. The datasets were then normalized to 10,000 counts per cell and log1p-transformed. The results section shows the OOD dataset (a) for which the models trained on scTab showed the highest overall performance.

#### NeurIPS Multiome Dataset

Our study included the NeurIPS multiome dataset^54^, a multi-modal bone marrow dataset that integrates gene expression counts with proteomics data. While distinct in its multi-omic nature, this dataset underwent similar preprocessing steps as our other datasets, ensuring consistency across all analyses. We split the dataset into train, validation, and test sets using an 80/10/10 random split. We chose 2,000 highly variable genes using Scanpy^61^ as a standard preprocessing step for this dataset.

### Self-Supervision Methods

#### Overview

Self-supervised learning (SSL) is the concept that data, along with its inherent pairwise relationships, are sufficient for learning meaningful data representations, even in the absence of explicit labels. While supervised learning relies on paired observations and labels (*X, Y*), self-supervised learning thus only depends on the input *X* and an inter-sample relationship (*X, G*), where *G* is constructed through a data augmentation that sustains the semantic information of *X*^8^. Thereby, the method distills signal from noise^62^, a crucial aspect for managing challenges like class imbalances in large, real-world datasets^63^. In single-cell data, this means distilling the signal of the cellular omics and removing noise sources such as batch effects or inconsistent labeling.

In the context of single-cell genomics (SCG), SSL harnesses these capabilities to navigate the complexities of vast, unlabeled datasets replete with intricate biological interdependencies. The framework is structured into two distinct phases: pre-training and fine-tuning. During the pre-training phase, the model employs contrastive learning or denoising methods to learn a data representation. This representation, characterized by its broad applicability, is then utilized in one of two ways. Firstly, as a ‘Zero-Shot SSL’ model, it can be directly applied to a downstream task without further label-dependent training. Alternatively, as an ‘SSL’ model, it undergoes fine-tuning to enhance performance on specific tasks. This fine-tuning capitalizes on the rich data representation acquired during pre-training, adjusting and optimizing it for the desired application. The fine-tuning phase of SSL, therefore, is not only about refining the pre-training but also about strategically leveraging the pre-established data mappings for task-specific optimizations.

#### Core Principles and Strategies

The choice of self-supervised pre-training, i.e., learning the inter-sample relationship, is critical to obtaining a meaningful data representation as it gives rise to the signal-to-noise distinction in the dataset. Our SSL framework is designed around two primary pre-training strategies: masked autoencoders and contrastive learning, both adapted to meet the unique demands of SCG.

Masked Autoencoders (MAEs): This approach follows the concept of self-prediction, where a significant portion of input features (genes in SCG) are masked (*i*.*e*., set to 0), and the model is trained to reconstruct these missing parts^9,42,64^. It thus sets focus on inter-feature dependencies. We implemented various masking strategies. (1) In Random Masking, 50% of genes are randomly chosen and masked with new choices in each iteration. (2) In Gene Program (GP) Masking, sets of genes known for biological functions are masked such that n% of genes are masked and reconstructed. The C8: cell type signature gene sets from the Human MSigDB Collections^65–67^ were used. Next, we introduce isolated Masked Autoencoders (iMAEs), in which all genes but a defined set are masked, and only this set is reconstructed. (3) For this, we present a gene program for transcription factor (GP to TF) isolated masking. This masking predicts the expression value of the transcription factor known to correspond to a gene program. This connection is given in the TFT: transcription factor targets subset of C3: regulatory target gene sets from the Human MSigDB Collections^68,69^. (4) Last, we present a gene program to gene program (GP to GP) isolated masking. In this strategy, a gene program is kept unmasked and used to predict only itself. The gene programs for this strategy also stem from the C8: cell type signature gene sets from the Human MSigDB Collections. These strategies are tailored to capture specific gene interactions and relationships, making them particularly suited for the intricate nature of single-cell data.

Contrastive Learning: Unlike self-prediction, contrastive learning focuses on understanding relationships between different samples, thus focusing on inter-sample dependencies. This method minimizes distances between similar samples and maximizes distances between dissimilar ones in the embedded space. Contrastive methods are typically distinguished by their strategy to avoid representation collapse, the trivial solution to contrastive losses of constant representations^9,41^. Bootstrap Your Own Latent (BYOL) is an example of architectural regularization through its teacher-student network. At the same time, Barlow Twins is an example of an information maximization method that avoids collapse by maximizing the information content of the embedding. We incorporated BYOL and Barlow Twins in our framework to benchmark two schools of thought. We used Gaussian noise for data augmentation, simulating the variable sequencing depth in RNA-seq experiments post-normalization and log1p transformation, thus reflecting typical noise profiles in SCG data.

#### Zero-Shot SSL Concept

A key concept in our study is the differentiation between the Zero-Shot SSL and SSL models. The Zero-Shot SSL model represents the initial phase of pre-training, where the model learns from the data without any label guidance through self-supervision algorithms. This model, even without fine-tuning, can provide meaningful insights into data, as demonstrated in various downstream tasks. The SSL model, on the other hand, undergoes an additional fine-tuning phase tailored to specific downstream applications. This distinction allows us to explore the full spectrum of SSL’s capabilities, from a generalized understanding of data to specialized, task-specific optimizations.

In summary, our self-supervision methods in SCG are defined by a nuanced application of masked autoencoders and contrastive learning adapted to the field’s specific challenges. The Zero-Shot SSL concept plays a central role in our approach, highlighting the potential of SSL to derive meaningful insights from large-scale, unlabeled datasets. This methodological framework sets the stage for a detailed exploration and benchmarking of SSL’s impact on various SCG tasks, as detailed in the following sections of our study.

### Downstream Applications in SCG

#### Cell Type Annotation

Cell type annotation in single-cell genomics is a classification task where data samples, represented as vectors of RNA-sequencing counts, are assigned to distinct cellular identities. Though seemingly straightforward, this task is complicated by the noise and heterogeneity inherent in large-scale datasets. We utilize the scTab dataset as the primary basis for our cell type annotation analysis. We employ various SSL methods and compare their effectiveness against supervised approaches. We train the classifier using a cross-entropy loss. We evaluate cell type annotation performance by k-nearest-neighbors (kNN, k=5) classification using the scTab validation set as neighbors of the test sample. This choice is driven to have the same evaluation, including for the Zero-Shot SSL model that does not have a prediction head. Our evaluation metrics focus on the macro F1 score, reflecting the models’ ability to handle class imbalances, supplemented by the micro F1 score, offering an additional comparative perspective to class imbalances.

List of used hyperparameters after parameter search:

| Parameter | Value |
| --- | --- |
| Hidden Layers | [512, 512, 256, 256, 64] |
| Batch Size | 16,384 |
| Dropout | 0.1 |
| Weight Decay | 0.05 |
| Learning Rate | 9e-04 |
| Learning Rate Scheduler | torch.optim.lr_scheduler.StepLR, gamma 0.9, step size 2 |
| Optimizer | torch.optim.AdamW |

#### Gene Expression Reconstruction

Gene expression reconstruction, the process of reconstructing counts from the transcriptome, still presents challenges due to the inherent noise and dispersion in RNA-seq data. The popular scVI model^36^ inspires our approach and diverges in its use of input data. While scVI uses raw counts as input and models them as a negative binomial distribution, our method employs normalized data for consistency with other downstream tasks. Nonetheless, similar to scVI, we predict the parameters of the negative binomial distribution. This strategy of modeling distribution parameters rather than direct RNA-seq count prediction enhanced reconstruction accuracy in our experiments. We opt for a non-variational, fully connected autoencoder framework consistent with our cell type prediction approach. Performance evaluation encompasses mean squared error (MSE) and uniform and weighted explained variance. We reported the weighted explained variance to best reflect the actual reconstruction efficacy, accounting for class imbalances. We include the MSE and uniform explained variance in our framework as supplementary evaluation, and they were used in our experiments.

List of used hyperparameters after parameter search:

| Parameter | Value |
| --- | --- |
| Hidden Layers | [512, 512, 256, 256, 64] |
| Batch Size | 8192 |
| Dropout | 0.05 |
| Weight Decay | 0.01 |
| Learning Rate | 0.001 |
| Learning Rate Scheduler | torch.optim.lr_scheduler.StepLR, gamma 0.9, step size 2 |
| Optimizer | torch.optim.AdamW |

#### Cross-Modality Prediction

Cross-modality prediction is the task of predicting one modality, in our case protein counts, from another, in our case RNA-sequencing counts. If performed well, such a task could augment cellular data by a different modality, offering a novel perspective. To test that, we use CITE-seq data from the NeurIPS multiome dataset. If fine-tuned, the models are trained to predict the protein modality from the RNA modality.

List of used hyperparameters after parameter search:

| Parameter | Value |
| --- | --- |
| Hidden Layers | [256, 256, 40] |
| Batch Size | 16384 |
| Dropout | 0.1574 |
| Weight Decay | 0.0012 |
| Learning Rate | 8.16e-05 |
| Learning Rate Scheduler | torch.optim.lr_scheduler.LambdaLR, step size 1 |
| Optimizer | torch.optim.Adam |

#### Data integration

Data integration is an effort to study a set of related single-cell genomics datasets, possibly curated from various donors with different pipelines and in different settings that create batch effects and technical artifacts. The scIB^60^ integration benchmarking is a well-established analysis to determine how well the relevant and meaningful biological signals are preserved in any model data representation while removing the unwanted batch effects resulting in a mixed representation of various datasets. Accordingly, the scIB pipeline measures two metrics, including the Bio Conservation and Batch Correction metrics, each consisting of several evaluations through different methods.

List of used hyperparameters after parameter search:

| Parameter | Value |
| --- | --- |
| Hidden Layers | [512, 512, 256, 256, 64] |
| Batch Size | 8192 |
| Dropout | 0.05 |
| Weight Decay | 0.01 |
| Learning Rate | 0.001 |
| Learning Rate Scheduler | torch.optim.lr_scheduler.StepLR, gamma 0.9, step size 2 |
| Optimizer | torch.optim.AdamW |

## Supporting information

Detailed Experimental Results

## Acknowledgments

We thank Felix Fischer for his valuable assistance with scTab and his constructive comments, which improved our work’s narrative. For the cross-modality prediction task, we thank Anastasia Litinetskaya for her valuable feedback. We also thank Merle Stahl, Xichen Wu, and Anna Chernysheva for their contributions during their master practical course, which laid the groundwork for this task (together with Yufan Xia, who continued afterward). We are particularly grateful to Alessandro Palma for his feedback on the manuscript’s storyline and to Alessandro Palma, Artur Szałata, and Eljas Roellin for their valuable input on the manuscript, greatly enhancing its quality. We thank Dr. Fabiola Curion for her input, sparking our exploration of isolated Masked Autoencoders, and her feedback on the multiomics application.

T.R. and M.B. are supported by the Helmholtz Association under the joint research school ‘Munich School For Data Science’. T.R. and F.J.T. acknowledge support by the Helmholtz Association’s Initiative and Networking Fund through CausalCellDynamics (grant # Interlabs-0029), F.J.T. acknowledges support by the European Union (ERC, DeepCell - 101054957). Views and opinions expressed are, however those of the author(s) only and do not necessarily reflect those of the European Union or the European Research Council. Neither the European Union nor the granting authority can be held responsible for them.

The language of this manuscript was refined using ChatGPT by OpenAI and Grammarly by Grammarly Inc.

## Author contributions

T.R., D.S.F., and F.J.T. conceptualized the project. T.R. led pilot analyses, method development, and implementation. T.R. and F.J.T. outlined the downstream analyses. T.R. performed the cell type prediction and gene expression reconstruction studies. T.R. and Y.X. undertook the cross-modality prediction analysis, and M.B. performed the data integration study. The manuscript was written by T.R., M.B., D.S.F., and F.J.T., with all authors contributing to discussions and providing comments on the manuscript.

## Code availability

The code is available at github.com/theislab/ssl_in_scg

## Data availability

The scTab data is available with instructions in the corresponding publication^4^. The smaller datasets are publicly available on CELLxGENE^48^ and subsets of the scTab datasets (HLCA: Dataset ID 148, PBMC: Dataset ID 87, Tabula Sapiens: Dataset ID 41). The novel datasets are sourced from CELLxGENE^48^ with instructions in the corresponding publications^49–52^. The NeurIPS multiome dataset is publicly available from NCBI GEO under accession GSE194122 with instructions in the corresponding publication^54^.

