## Supplementary material for "Delineating the Effective Use of Self-Supervised Learning in Single-Cell Genomics": Detailed Experimental Results

### Appendix

#### Supp. Figure 1: Detailed Results of the Evaluation on Individual Datasets

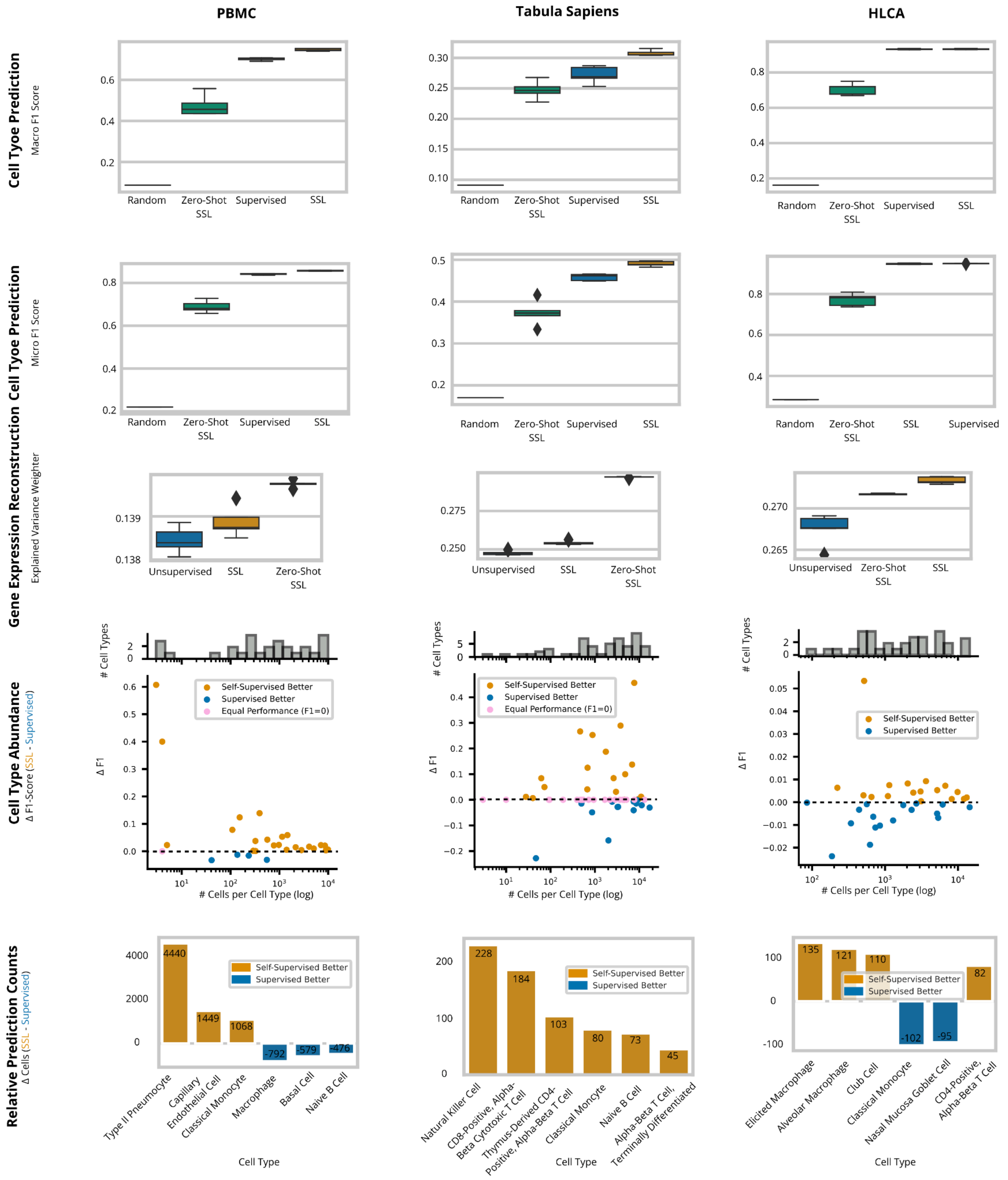


**Supp Figure 2: Self-supervised learning improves downstream performance on PBMC, Tabula Sapiens, and HLCA datasets**

Columns show the three datasets or atlases involved in this evaluation. The first row shows box plots of Macro F1 scores for the cell-type prediction performance of (i) the random baseline, (ii) the Zero-Shot SSL model, (iii) the supervised model, and (iv) the SSL model. The second row supplements the cell-type prediction performance with the Micro F1 score. The third row shows the gene expression prediction performance of (i) the Zero-Shot SSL model, (ii) the supervised model, and (iii) the SSL model using the weighted explained variance. The fourth row shows the relative performance of the SSL model to the supervised model at the hand of the Macro F1 difference. The points represent a cell type, and the relative performance is plotted against the abundance of that cell type as a log number of cells per cell type. Additionally, a bar plot indicates how many cell types of an abundance exist in the dataset. The fifth row shows the relative number of correctly predicted cells per cell type for the 5 cell types with the largest absolute difference.

#### Supp. Figure 2: Detailed Results of the Evaluation of Novel Datasets

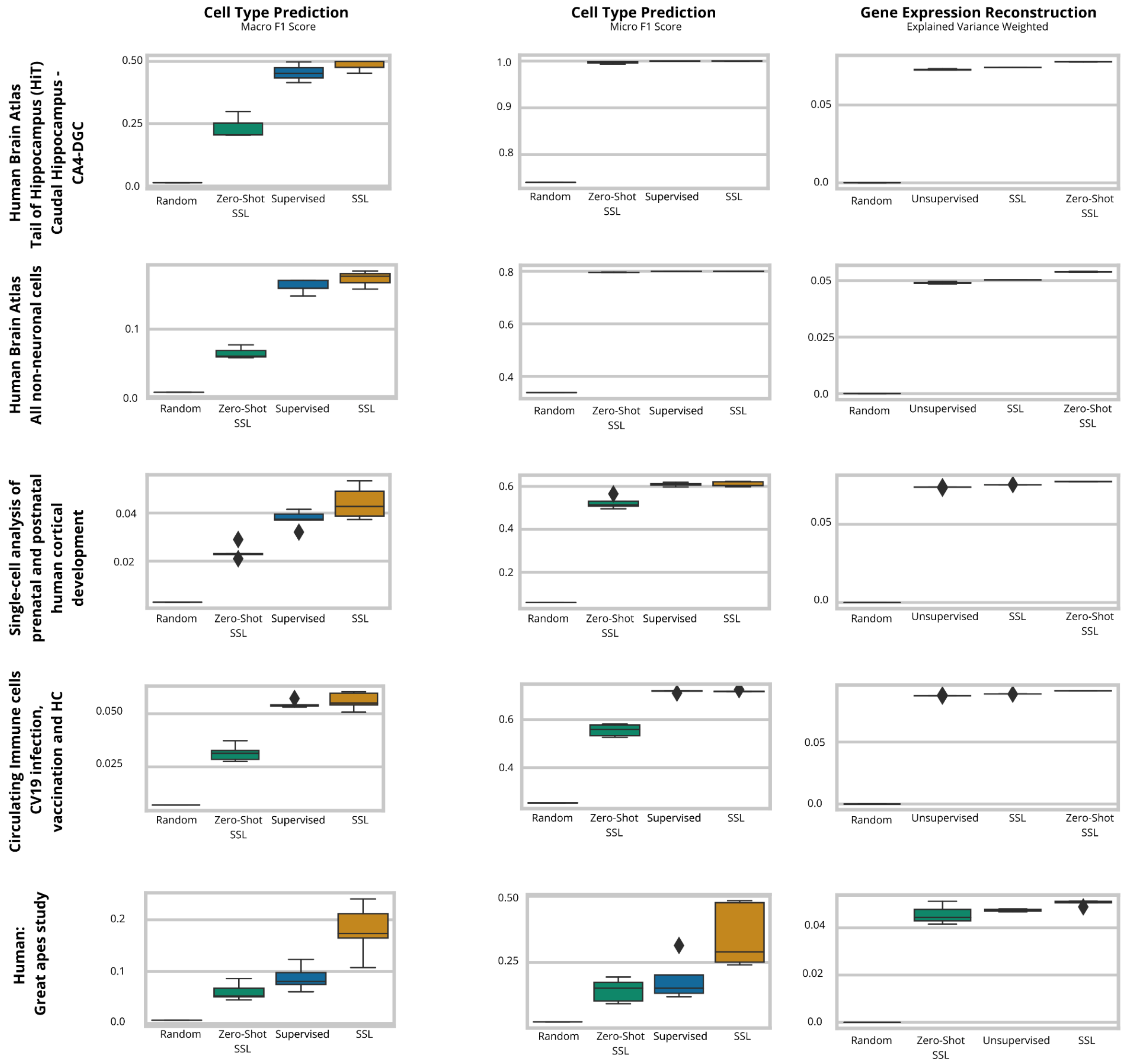


**Supp Figure 3: Self-supervised learning offers advantages when analyzing novel datasets**

Columns show the task of cell type prediction, evaluated with the Macro F1 (left column) and the Micro F1 (middle column) score, and the gene expression reconstruction task, evaluated with the weighted explained variance. The columns show the five novel datasets that are not included in the scTab dataset.
